# Generalized fear following acute stress is caused by change in co-transmitter identity of serotonergic neurons

**DOI:** 10.1101/2023.05.10.540268

**Authors:** Hui-quan Li, Wuji Jiang, Lily Ling, Vaidehi Gupta, Cong Chen, Marta Pratelli, Swetha K. Godavarthi, Nicholas C. Spitzer

**Affiliations:** Neurobiology Department and Kavli Institute for Brain and Mind, University of California, San Diego; Department of Cellular and Molecular Medicine, University of California, San Diego

## Abstract

Overgeneralization of fear to harmless situations is a core feature of anxiety disorders resulting from acute stress, yet the mechanisms by which fear becomes generalized are poorly understood. Here we show that generalized fear in mice in response to footshock results from a transmitter switch from glutamate to GABA in serotonergic neurons of the lateral wings of the dorsal raphe. We observe a similar change in transmitter identity in the postmortem brains of PTSD patients. Overriding the transmitter switch in mice using viral tools prevents the acquisition of generalized fear. Corticosterone release and activation of glucocorticoid receptors trigger the switch, and prompt antidepressant treatment blocks the co-transmitter switch and generalized fear. Our results provide new understanding of the plasticity involved in fear generalization.

**One sentence summary:** Acute stress produces generalized fear by causing serotonergic neurons to switch their co-transmitter from glutamate to GABA

## Introduction

Fear, when properly controlled, is an essential emotion for survival, because it enables risk evaluation and promotes vigilance (*1*). Acquired fear responses can be specific to the cue or context associated with an aversive stimulus (defined as conditioned fear), but often generalize to other environmental cues or contexts (defined as generalized fear) (*2*). Fear generalization can be harmful when fear responses are sustained and inappropriate to environmental stimuli.

Although fear generalization occurs in many stress- and fear-related disorders (*3*), the cellular and circuit mechanisms by which fear is generalized remain largely unknown (*2*). Behavioral correlates of generalized fear are regulated by the activity of neurotransmitters within neural circuits involving the ventrolateral periaqueductal gray (vlPAG) and the dorsal raphe nucleus (DRN) (*4*). The vlPAG and DRN overlap in the lateral wings of dorsal raphe (lwDR) (*5, 6*). Activity in serotonergic neurons in the lwDR (5-HT^lwDR^) has been linked with panic-like fear responses (*7, 8*).

The ability of 5-HT^lwDR^ neurons to co-release GABA, glutamate and serotonin complicates study of the behavior of this population of neurons in neural circuits (*9–11*). We examined each of these neurotransmitter phenotypes and determined how they change in response to stress that evokes fear. Here we identify adult brain plasticity in which 5-HT^lwDR^ neurons change their co-transmitter in response to acute stress and cause persistent generalized fear.

## Results

### Footshock induces generalized fear correlated with a glutamate-to-GABA switch in the lwDR

To understand the mechanisms underlying generalization of fear responses, we studied contextual fear. Pairing of an unconditioned stimulus (US; footshock) and the conditioning context (CS) leads to conditioned fear, when testing occurs later in the same context (A) (*2, 12, 13*). Increasing the intensity of the US produces fear generalization even when testing occurs later in a new context (B), with different shape, size, texture and color (*13–15*). We validated these findings by testing the effects of a single weak footshock (0.5 mA, 1 s) and 5 strong (1.5 mA, 2 s) footshocks on acquisition of conditioned and generalized fear in the mouse. Two weeks later we measured freezing behavior in the same context (A) and in a different context (B). We calculated a freezing score as time spent freezing)/(total time in the context)x100%. The weak shock led to enhanced freezing in the training context (A) (shock 55% vs. no shock 4%; **Fig. 1a,b**) but not in the new context (B) (shock 4% vs. no shock 3%; **Fig. 1c**). Notably, strong shock led to more freezing in both context A (shock 64% vs. no shock 3%; **Fig. 1b**) and context B (shock 29% vs. no shock 2%; **Fig. 1c**). We confirmed this difference in freezing after weak shock or strong shock in context C, which had features different from those of either context A or B (strong shock 27% vs no shock 3%; weak shock 4% vs no shock 3%) (**Supplementary Fig. 1a,b**). Exploration behavior in contexts B and C also decreased after strong shock (distance traveled and amount of rearing) (**Fig. 1d,e; Supplementary Fig. 1c,d**). In addition, the number of fecal boli produced in these two contexts increased (**Fig. 1f**; **Supplementary Fig. 1e**). These results demonstrate that although both weak shock and strong footshock were sufficient to cause conditioned fear, only the strong footshock triggered generalized fear.

**Fig. 1.**
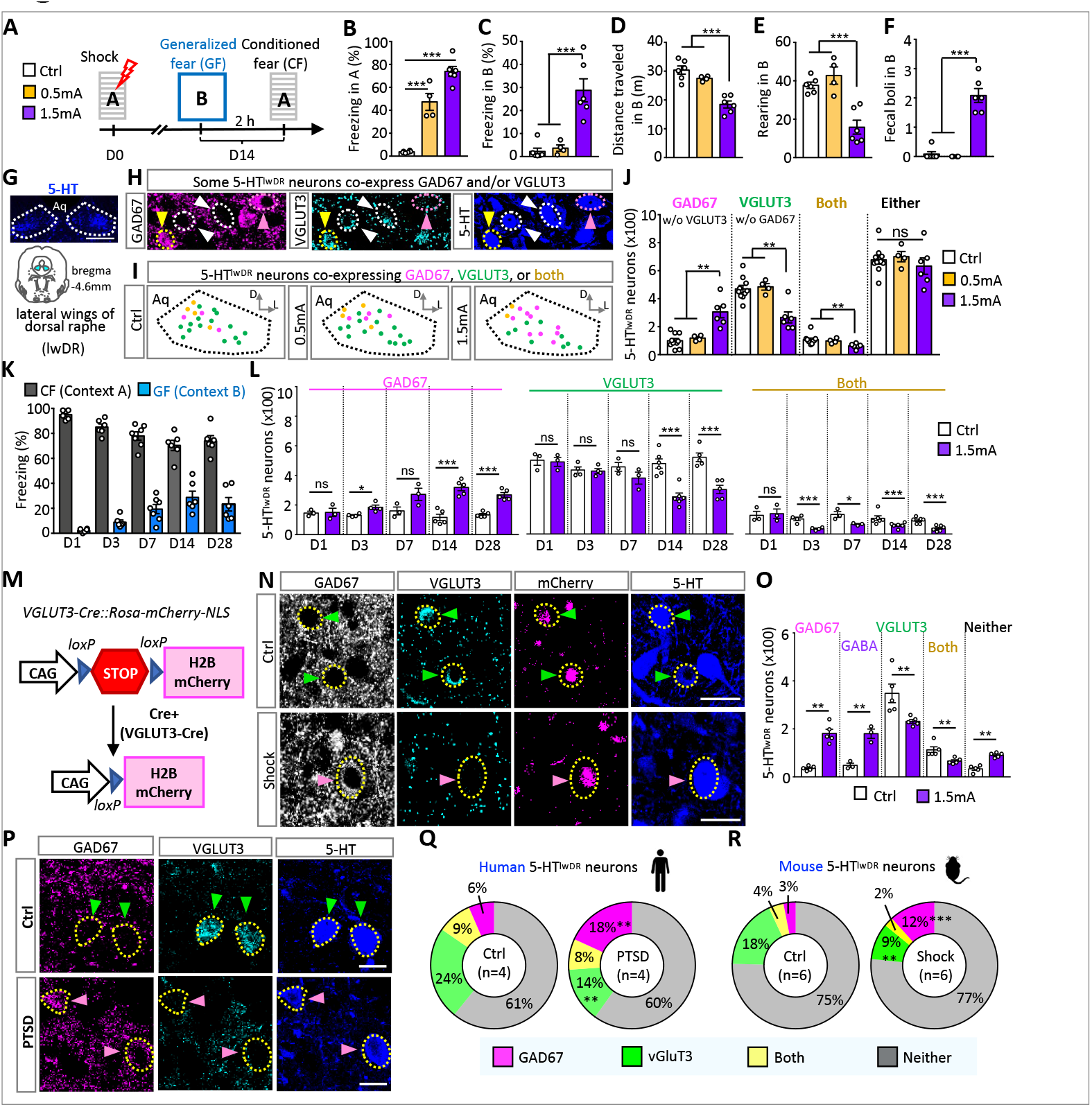
Strong footshock induces generalized fear and a switch from VGLUT3 to GAD67 in serotonergic neurons. (**A**) Experimental design. Mice were given no shock, a single shock (0.5 mA, 1 s), or 5 shocks (1.5 mA, 2 s) separated by 40 to 60 s in context A. Mice were later tested for fear responses in context B and 2 hr later in context A. (**B**, **C**) Freezing in contexts A and B. (**D, E**) Horizontal and vertical exploration; (**F**) Number of fecal boli in context B. (**B**-**F**) n=4-6 mice/group. (**G**) Upper: staining of 5-HT (serotonin) in the dorsal raphe. Dotted lines outline the lateral wings. Lower: coronal section of mouse brain. Light blue indicates the lwDR. (**H**) Triple staining of the lwDR. Pink arrowheads, 5-HT neurons expressing GAD67 but not VGLUT3. Cyan arrowheads, 5-HT neurons expressing VGLUT3 but not GAD67. Brown arrowheads, 5-HT neurons co-expressing both GAD67 and VGLUT3. (**I**) Location of 5-HT neurons expressing VGLUT3 (green), GAD67 (pink), and both (brown) in coronal sections of mice given shock or no shock. (**J**) Number of each neuronal phenotype 2 weeks after shock. n=4-8 mice/group. (**K**) Development of fear (day 1-28) in contexts A and B. n=3-8 mice/group. (**L**) Development of changes in transmitter phenotype. n=3-6 mice/group. (**M**) Experimental design to permanently label VGLUT3 neurons with mCherry. (**N**) Quadruple-staining of the lwDR in VGLUT3-Cre::Rosa-mCherry-NLS mice 2 weeks after shock or no shock control. Green arrowheads, 5-HT neurons expressing mCherry, VGLUT3 but not GAD67. Pink arrowheads, 5-HT neurons expressing mCherry, GAD67 but not VGLUT3. (**O**) Neuron numbers in VGLUT3-Cre::Rosa-mCherry-NLS mice 2 weeks after shock. n=3-5 mice/group. (**P**) Triple staining of the lwDR in human brains. (**Q**) Percent of 5-HT^lwDR^ neurons expressing each transmitter marker. Human, n=1269 cells for Ctrl, 1232 cells for PTSD; n=4 patients/group. Mice, n=2206 cells for no shock, 2177 cells for shock; n=6 animals/group.

To investigate whether weak or strong footshock causes neurotransmitter switching, we immunostained the dorsal raphe two weeks after strong shock (**Fig. 1g-i**), at a time when generalized fear was observed (**Fig. 1a,c**). We quantified the numbers of neurons in the lateral wings (lwDR) that co-express 5-HT, GAD67 (glutamate decarboxylase 67, a synthetic enzyme for GABA) and VGLUT3 (a vesicular glutamate transporter). Weak shock did not change the number of 5-HT^lwDR^ neurons or the number of these neurons co-expressing either GAD67 or VGLUT3 (**Fig. 1j**). However, strong shock increased the number of 5-HT^lwDR^ neurons that co-express GAD67 and not VGLUT3 (shock, 305±41; no shock, 93±25) and decreased the number of 5-HT^lwDR^ neurons co-expressing VGLUT3 and not GAD67 (shock, 266±39; no shock, 504±33). 5-HT^lwDR^ neurons that co-express both GAD67 and VGLUT3 also decreased in mice receiving strong shock (shock, 62±7; no shock, 113±18) (**Fig. 1j**). The total number of 5-HT^lwDR^ neurons that co-express either GAD67 or VGLUT3 showed no change (shock, 633±59; no shock, 710±37). The total number of 5-HT^lwDR^ neurons overall, independent of the presence or absence of co-transmitters, was also unchanged (shock, 2454±148; no shock, 2293±71) (**Supplementary Fig. 2a**).

We next examined the change in the transmitters themselves by staining GABA and glutamate in the lwDR. Consistent with the increase in the number of 5-HT^lwDR^ neurons that co-express GAD67 two weeks after strong shock (**Fig. 1j**), we found an increase in the number of 5-HT^lwDR^ neurons that co-express GABA (**Supplementary Fig. 2b-d**). These neurons also expressed glutamate, consistent with its role as a precursor for GABA (>99% of 5-HT^lwDR^ neurons that expressed GABA were immunoreactive for glutamate). Accordingly, the number of neurons that contain glutamate would be expected to be similar to the total number of neurons co-expressing either GAD67 or VGLUT3 (**Fig. S2d**). The results suggest that strong but not weak footshock triggers a glutamate-to-GABA switch in the 5-HT^lwDR^ (**Fig. 1g-j**; **Supplementary Fig. 2**). Both the transmitter switch and generalized fear were evoked by strong but not weak footshock, and weak shock was sufficient to cause only conditioned fear (**Fig. 1b,c**).

To determine the time course of the change in transmitter identity and the onset of conditioned and generalized fear, we measured freezing and examined VGLUT3 and GAD67 expression in the 5-HT^lwDR^ at 1, 3, 7 days, and 4 weeks after strong footshock. Conditioned fear was present 1 day after shock, but generalized fear was first observed at 3 days (**Fig. 1k**). Similarly, the change in neurotransmitter identity was not observed at 1 day after shock but was detected after 3 days and to an even greater degree after 1 week (**Fig. 1l**). Both the VGLUT3-to-GAD67 switch and generalized fear were sustained for at least 4 weeks (**Fig. 1l**). These results document a temporal correlation between the change in transmitter identity and the acquisition of generalized fear.

The absence of evidence for apoptosis or neurogenesis (**Supplementary Fig. 3a-e**) suggested that the increase in number of 5-HT^lwDR^ neurons co-expressing GAD67 was derived from neurons that had previously co-expressed VGLUT3. To seek direct evidence for this VGLUT3-to-GAD67 switch at the single-cell level, we selectively tagged glutamatergic neurons with a genetic marker to track changes in transmitter identity after footshock. We used a VGLUT3-Cre transgenic mouse line (*16*), in which a Cre recombinase is expressed under the VGLUT3 promoter, crossed with a Rosa26-CAG-LSL-H2B-mCherry (referred to as Rosa-H2B-mCherry) mouse line with a loxP-flanked STOP sequence followed by a nuclear-targeted H2B-mCherry fusion protein. In these VGLUT3-Cre::Rosa-H2B-mCherry doubly transgenic mice, VGLUT3-driven Cre induces cleavage of the STOP sequence, allowing continuous expression of mCherry in cell nuclei (**Fig. 1m**). As a result, the nuclei of neurons expressing VGLUT3 are permanently labeled with mCherry even if their expression of VGLUT3 is transient. We immunostained 5-HT, GAD67 and VGLUT3 to count the number of 5-HT+mCherry+ neurons that co-express each transmitter marker (**Fig. 1n**). We found that the number of 5-HT^lwDR^ mCherry neurons co-expressing only VGLUT3 but not GAD67 decreased and the number of 5-HT^lwDR^ mCherry neurons co-expressing GAD67 but not VGLUT3 increased (**Fig. 1o**). This result is consistent with a switch from VGLUT3 to GAD67 in single neurons after strong shock. The number of neurons co-expressing both GAD67 and VGLUT3 was smaller in 5-HT^lwDR^ mCherry neurons mice receiving strong shock than in mice receiving no shock (**Fig. 1o**), as it was in wildtype mice (**Fig. 1j**). As noted above, >99% of 5-HT^lwDR^ neurons that contain GABA are also immunoreactive for glutamate (**Supplementary Fig. 2c**). Accordingly, we quantified GABA and VGLUT3 (not glutamate) in the 5-HT^lwDR^ of VGLUT3-Cre::Rosa-H2B-mCherry mice and found that the number of neurons co-expressing GABA but not VGLUT3 increased after strong shock (**Fig. 1o**).

To understand whether fear responses and the VGLUT3-to-GAD67 switch were specific to males or occurred in both males and females, we gave strong shock to female VGLUT3-Cre::mCherry-NLS mice, tested their fear responses two weeks later, and examined transmitter identity in the 5-HT^lwDR^. Strong foostshock caused a VGLUT3-to-GAD67 switch in 5-HT^lwDR^ neurons and induced both conditioned and generalized fear (**Supplementary Fig. 4**).

Fear generalization occurs in patients with stress and fear disorders such as post-traumatic stress disorder (PTSD) who have experienced intense traumatic events (*15*). We examined transmitter identity in postmortem brains of four patients with PTSD and four control subjects of comparable age and post-mortem interval (**Supplementary Fig. 5**) (tissue provided by the NIH Neurobiobank). Patients, in comparison to controls, exhibited a decrease in the number of 5-HT^lwDR^ neurons co-expressing VGLUT3 and an increase in the number of neurons co-expressing GAD67 (**Fig. 1p-r**). The PTSD diagnosis was current at time of death and symptoms, including generalized fear, had persisted for at least 3 years. The altered expression of VGLUT3 and GAD67 in these patients matched the change in transmitter identity described here in mice that were given strong footshock and exhibited generalized fear (**Fig. 1c,h,j**).

### Appearance of generalized fear requires the glutamate-to-GABA switch in the 5-HT^lwDR^

To determine whether the VGLUT3-to-GAD67 switch is essential for the generalization of fear responses, we first investigated the functional relevance of the increase in number of neurons expressing GAD67. We bilaterally injected AAV-CAG-DIO-shGAD1-EGFP (*17*) into the lwDR of SERT-Cre mice to drive expression of GAD1 shRNA in serotonergic neurons, thereby interfering with the gain of neurons expressing GAD67 (**Fig. 2a**). AAV-CAG-DIO-shScr (scramble shRNA)-EGFP served as a control. Suppression of GAD67 expression by AAV-DIO-shGAD1-EGFP in SERT-IRES-Cre mice was confirmed by the absence of co-expression of EGFP with GAD67 (**Supplementary Fig. 6a,b**). Four weeks after AAV injection, mice were given strong footshock and tested for fear responses two weeks later (**Fig. 2b,c**). The mice that received shGAD1 followed by strong footshock (hereafter, shGAD1-shock) showed partially reduced freezing in context A compared to shScr-shock mice (from 59% to 38%) (**Fig. 2b**). Importantly, freezing in context B was wholly blocked by shGAD1 expression (from 21% to 3%)(**Fig. 2c**). Consistent with reduced freezing, shGAD1-shock mice also showed enhanced exploration activity (distance traveled, from 18 to 30 meters; rearing counts, from 17 to 38) (**Fig. 2d,e**) and reduced fecal boli (from 2 to 0)(**Fig. 2f**). These results suggest that the footshock-induced gain of GAD67 in 5-HT^lwDR^ neurons is necessary to produce generalized fear.

**Fig. 2.**
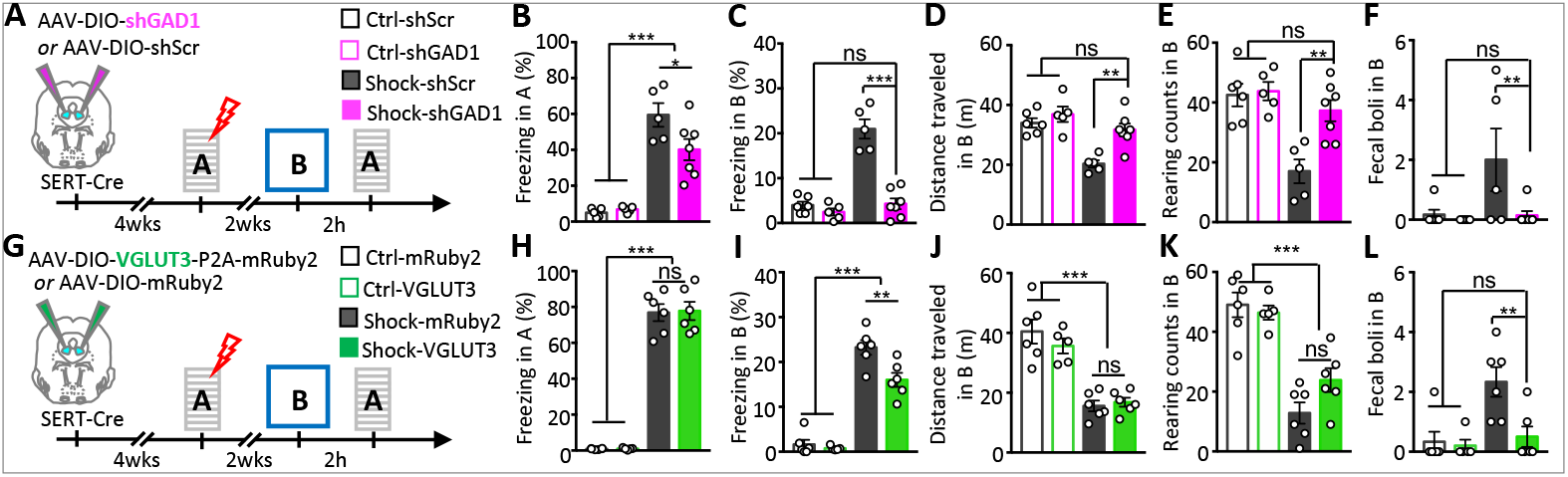
Blocking the glutamate-to-GABA switch in the lwDR suppresses the appearance of generalized fear. (**A**) Experimental design to override the gain of GAD67 in SERT-Cre neurons and assess behavior. (**B, C**) Freezing in contexts A and B for SERT-Cre mice that express AAV-DIO-shScr or AAV-DIO-shGAD1 in the lwDR. (**D-F**) Exploration and fecal production in context B. (**B-F**) n=5-7 mice/group. (**G**) Experimental design to replace the loss of VGLUT3 in SERT-Cre neurons and assess behavior. (**H, I**) Freezing in contexts A and B for SERT-Cre mice that express AAV-DIO-VGLUT3-mRuby2 or AAV-DIO-mRuby2 in the lwDR. (**J-L**) Exploration and fecal production in context B. (**H-L**) n=5-6 mice/group.

To test whether the loss of VGLUT3 is also essential for the generalization of fear, we generated AAV-hSyn-DIO-VGLUT3-P2A-mRuby2 (shortened as AAV-DIO-VGLUT3) that expresses Cre-dependent VGLUT3 driven by the human Synapsin (hSyn) promoter. Co-expression of viral mRuby2 with VGLUT3 validated Cre-dependent expression of VGLUT3 by AAV-DIO-VGLUT3 in SERT-IRES-Cre mice (**Supplementary Fig. 6a,c**). AAV-DIO-mRuby2 served as a control (**Fig. 2g**). We bilaterally injected the AAVs into the lwDR of SERT-IRES-Cre mice, allowed four weeks for AAV expression, then gave mice strong footshock and tested fear responses two weeks later (**Fig. 2h,i**). We found that VGLUT3-expressing mice given strong shock showed no effect on conditioned fear in context A and performed as mice expressing AAV-DIO-mRuby2 (78%) (**Fig. 2h**). However, fear responses in context B were reduced, as indicated by reduced freezing (from 23% to 15%) (**Fig. 2i**) and reduced number of fecal boli (from 2.3 to 0.4) (**Fig. 2l**). Exploration behavior was marginally increased (distance traveled, from 16 to 17 m; rearing counts, from 10 to 23) (**Fig. 2j,k**). These results imply that the footshock-induced loss of VGLUT3 in some 5-HT^lwDR^ neurons makes a smaller contribution to generalized fear responses than the gain of GAD67.

In addition to fear responses, mice exhibited an increased startle response and decreased prepulse inhibition following strong footshock (**Supplementary Fig. 7a,b**), similar to those reported for stressed rodents and human volunteers (*18–20*). However, the effect of shock on these behaviors was not affected by either knockdown of GAD67 expression or exogenous expression of VGLUT3 in 5-HT^lwDR^ neurons (**Supplementary Fig. 7c-f**). These results suggest that the VGLUT3-to-GAD67 switch in 5-HT^lwDR^ neurons induced by strong shock does not contribute to the effects on arousal or sensorimotor gating caused by stress.

### Switching neurons projecting to the central amygdala and lateral hypothalamus regulate generalized fear

5-HT^lwDR^ neurons project to nuclei regulating fear, including the central amygdala (CeA), lateral habenula (LHB), paraventricular nucleus of the hypothalamus (PVN) and lateral hypothalamus (LH) (*9, 21*). To learn which of these regions are innervated by the glutamatergic neurons that have acquired a GABAergic phenotype, we first injected a Cre-dependent anterograde tracer, AAV-hSyn-DIO-TdTomato-T2A-synaptophysin-EGFP (*17, 22*) into the lwDR of SERT-Cre mice to drive expression of TdTomato and SypEGFP in 5-HT^lwDR^ neurons. A T2A self-cleaving sequence located between TdTomato and SypEGFP separates the two markers, so that TdTomato labels cell bodies and axons, and the SypEGFP fusion protein labels only presynaptic terminals (**Fig. 3a**). After four weeks we gave mice strong footshock. Two weeks later we confirmed the correct expression of TdTomato in 5-HT^lwDR^ neuronal cell bodies (**Fig. 3b**) and then stained EGFP in the CeA, LHB, PVN and LH to identify putative synaptic terminals. We co-stained VGLUT3 or VGAT (vesicular GABA transporter) to trace presynaptic terminals that transport glutamate or GABA into synaptic vesicles (**Fig. 3c,d**). The percent of putative EGFP+ synaptic terminals labeled with VGLUT3 or VGAT was quantified by calculating Manders’ colocalization coefficients. These experiments demonstrated a decrease in the percent of EGFP overlapped with VGLUT3 and an increase in the percent of EGFP overlapped with VGAT, specifically in the LH and CeA but not in the LHB or PVN of shocked mice compared to no-shock controls (**Fig. 3e,f**). These results suggested that 5-HT^lwDR^ neurons projecting to the LH and CeA switch the expression of VGLUT3 to VGAT at their axon terminals.

**Fig. 3.**
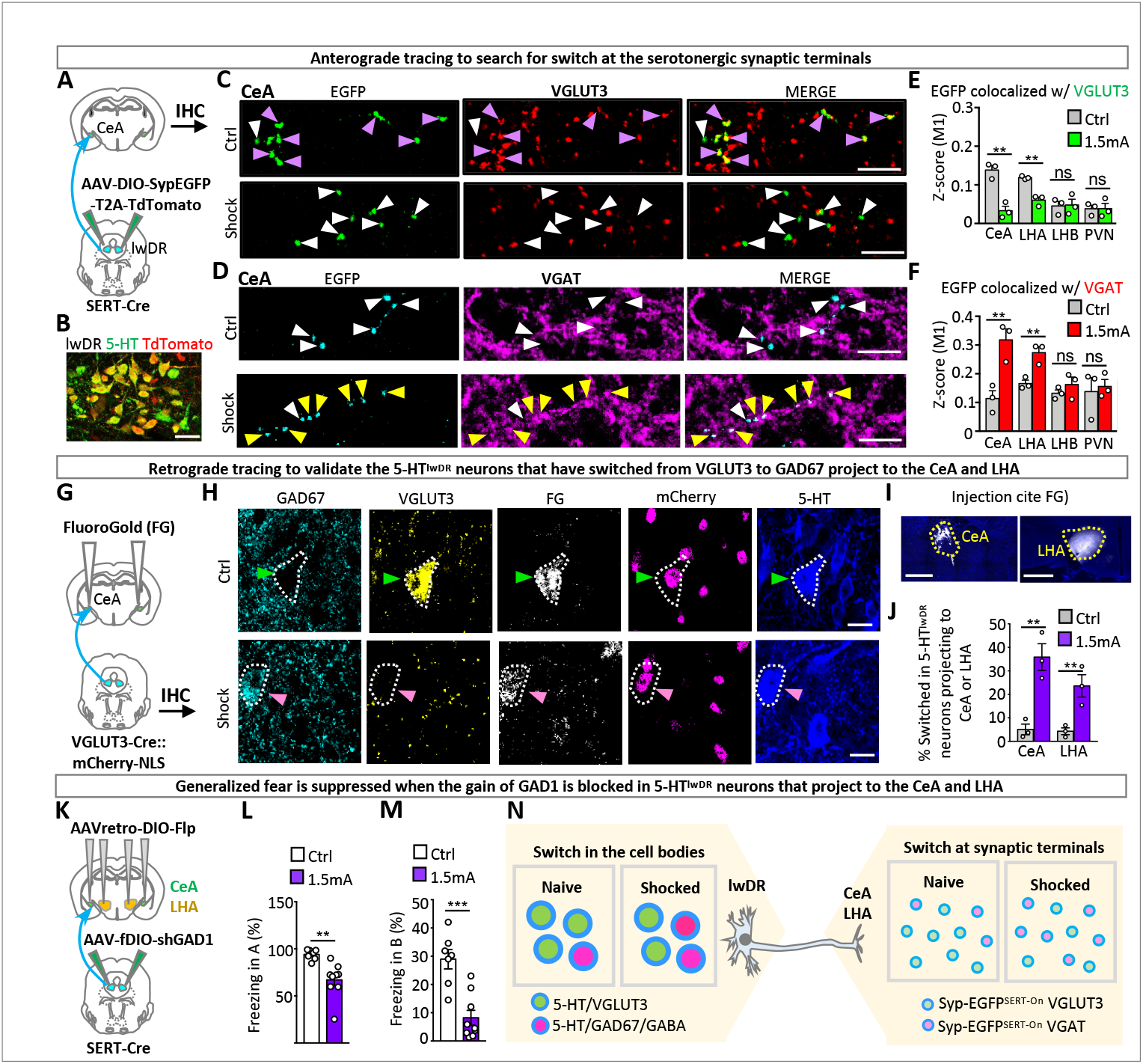
Switching neurons regulating generalized fear project to the central amygdala and lateral hypothalamus. (**A**) Experimental design to identify presynaptic changes in transporter expression of serotonergic lwDR axons. AAV8-phSyn1-FLEX-tdTomato-T2A-Syptophysin-EGFP-WPRE (AAV-FLEX-TdTomato-T2A-SypEGFP) was injected into the lwDR of SERT-Cre mice. Coronal sections containing the CeA, LHA, LHB or PVN were immunostained for EGFP and VGLUT3 or EGFP and VGAT. (**B**) Double immunostaining of TdTomato and 5-HT in the lwDR of mice that received 1.5 mA shock. (**C, D**) Double immunostaining of VGLUT3 or VGAT and EGFP in the CeA of shocked or control mice injected with AAV-FLEX-SypEGFP-T2A-TdTomato in the lwDR. Scale bar, 10 μm. (**E, F**) The M1 coefficient of VGLUT3 or VGAT colocalization with putative serotonergic presynaptic terminals. N>800 EGFP+ terminals/mouse; n=3 mice/group. (**G**) Experimental design to validate the targets of 5-HT^lwDR^ neurons that have switched from co-expressing VGLUT3 to coexpressing GAD67. (**H**) Quintuple-labeling of GAD67, VGLUT3, fluorogold, mCherry and 5-HT in the lwDR of both control and shocked mice. Scale bar, 10 μm. (**I**) Representative coronal sections show fluorogold (white) injected into the CeA or LHA. Scale bar, 1 mm. (**J**) Percent of neurons expressing 5-HT, mCherry and fluorogold that express GAD67 but not VGLUT3. The x-axis indicates the brain regions in which fluorogold was injected. n=3 mice/group. (**K**) Experimental design to test whether blocking the gain of GAD67 in 5-HT^lwDR^ neurons that project to the CeA or LHA suppresses generalized fear. (**L**) Conditioned and (**M**) generalized fear of mice injected in (**K**), housed for 4 weeks, shocked at 1.5 mA and tested two weeks later. n=5-8 mice/group. (**N**) Cartoon showing the switch from VGLUT3 to GAD67 in the serotonergic cell bodies is consistent with a switch in transporter expression from VGLUT3 to VGAT at serotonergic terminals in the CeA and LHA.

We next investigated whether the 5-HT^lwDR^ neurons that have switched from VGLUT3 to GAD67 in their cell bodies project to the CeA and LHA. To do so, we combined retrograde tracing with the genetic approach used to identify the switch from VGLUT3 to GAD67 in single neurons (see **Fig. 1m-o**). We bilaterally injected the fluorescent retrograde axonal tracer FluoroGold into the LH or CeA of VGLUT3-Cre::Rosa-H2B-mCherry mice (**Fig. 3g**). Neurons that project to the LH or CeA were labeled with FluoroGold, and neurons that expressed VGLUT3 were labeled with mCherry. One-week later mice were given strong shock and sacrificed for immunostaining two weeks later. When FluoroGold was injected into the LH (or CeA), lwDR cells that contained FluoroGold and 5-HT were identified as 5-HT^lwDR->LH^ (or 5-HT^lwDR->CeA^) neurons. Some 5-HT^lwDR->LH^ (or 5-HT^lwDR->CeA^) cells were positive for mCherry and GAD67, but not for VGLUT3. These cells were recognized as neurons that project to the LH (or CeA) and have switched expression of VGLUT3 to GAD67 (**Fig. 3h,i)**. We found an increase in the number of these neurons in shocked compared to control mice when Fluorogold was injected either in the LH or in the CeA (**Fig. 3j)**, demonstrating that newly GABAergic neurons project to both nuclei.

We then determined whether suppression of the GABAergic phenotype only in 5-HT^lwDR->LH^ or 5-HT^lwDR->CeA^ neurons was sufficient to block generalized fear. To do this, we injected a Cre-dependent retrograde AAV expressing a Flp recombinase (AAVretro-DIO-Flp) into the LH (or CeA) of SERT-Cre mice and co-injected a Flp-dependent AAV expressing shRNA for GAD1 (AAV-fDIO-shGAD1) into the lwDR (**Fig. 3k**). This protocol enabled expression of shRNA for GAD1 specifically in 5-HT^lwDR^ neurons that project to the LH or to the CeA. Mice injected with AAVretro-DIO-Flp and AAV-fDIO-shScr served as controls. We found that shGAD1 expression in both 5-HT^lwDR->LH^ and in 5-HT^lwDR->CeA^ neurons had a mild effect on conditioned fear (**Fig. 3l**) but significantly blocked generalized fear (**Fig. 3m,n**).

### Activation of the CRH/CORT/GR signaling pathway causes the glutamate-to-GABA switch

We next investigated how footshock generates the VGLUT3-to-GAD67 switch. Footshock is a stressor for rodents, activating the hypothalamic-pituitary-adrenal (HPA) axis that is the principal organizer of the body’s response to stress. The HPA axis comprises corticotropin-releasing hormone (CRH) secreted by the PVN, adrenocorticotropic hormone (ACTH) released by the anterior pituitary, and corticosterone (CORT) released by the adrenal cortex, activating their cognate receptors (*23*). The dorsal raphe expresses glucocorticoid receptors, GR, that modulate fear and anxiety (*11, 24, 25*). Knockout of GR affects the neurotransmitter phenotype in the dorsal raphe (*26*). These findings led us to hypothesize that the HPA axis plays a role in the generation of the VGLUT3-to-GAD67 switch in the dorsal raphe (**Fig. 4a**).

**Fig. 4.**
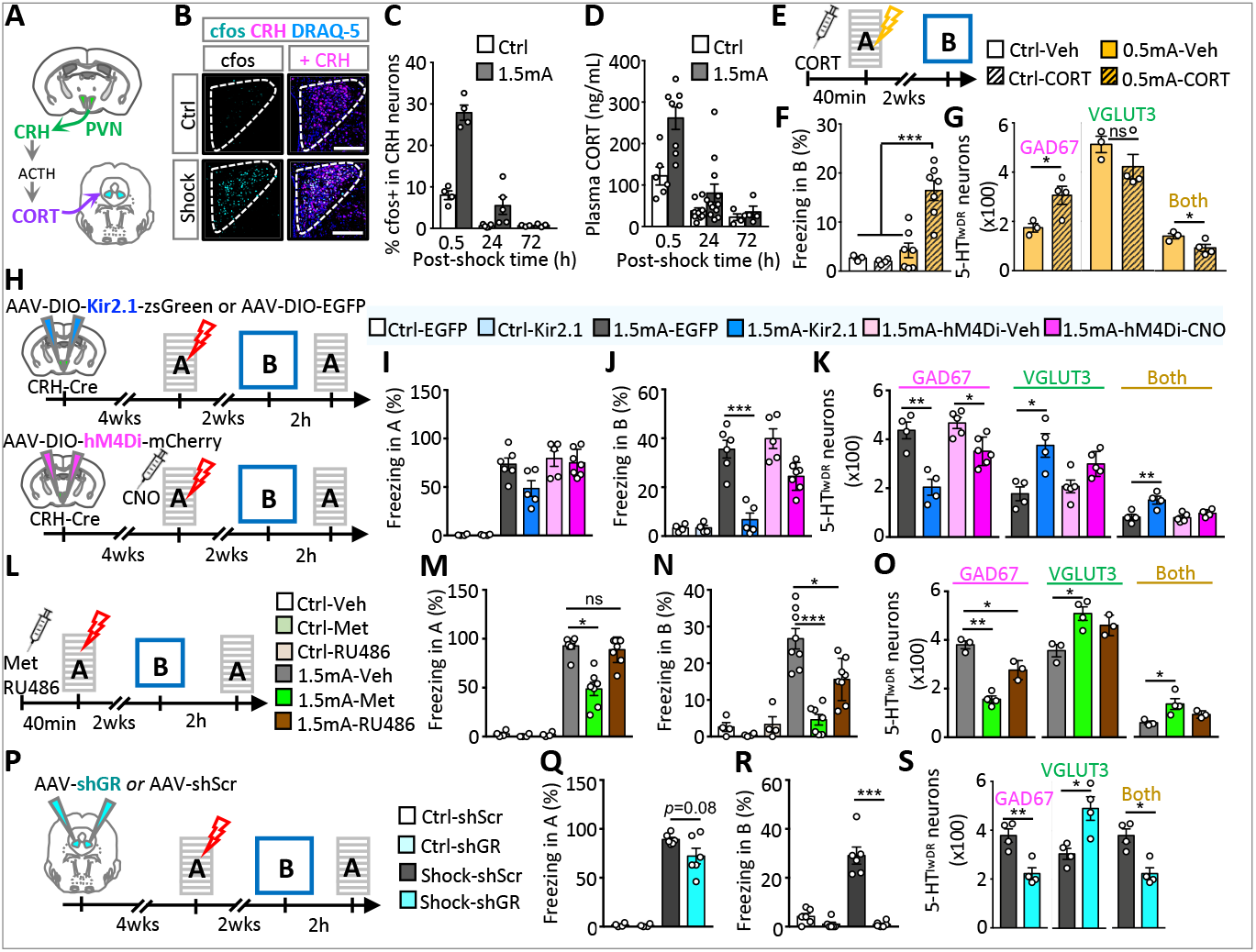
Activation of the CRH/CORT/GR signaling pathway causes the switch in serotonergic neurons and generalized fear. (**A**) Schematic of the hypothalamic-pituitary-adrenal axis. (**B**) Triple staining of the PVN in mice sacrificed 30 min post-shock and in no-shock controls. (**C**) Time course for c-fos expression in CRH^PVN^ neurons; n=100 to 150 cells/mouse; n=4 mice/group. (**D**) Time course for the level of plasma CORT; n=6-12 mice/group. (**E**) Experimental design to test whether CORT increases generalized fear. (**F**) Generalized fear of mice injected with CORT or Veh, given a single 0.5 mA shock or no shock. (**G**) Numbers of neurons coexpressing GAD67 or VGLUT3 in mice sacrificed immediately after behavior tests. n=4 mice/group. (**H**) Experimental design to block activity of CRH^PVN^ neurons and test fear. (**I, J**) Conditioned and generalized fear of CRH-Cre mice injected with AAV-DIO-Kir2.1-zsGreen, AAV-DIO-EGFP, or AAV-DIO-hM4D(Gi). n=5-7 mice/group. (**K**) Numbers of neurons coexpressing GAD67 or VGLUT3 in mice sacrificed immediately after behavior tests. n=4-5 mice/group. (**L**) Experimental design to block CORT synthesis by injecting Metyrapone (Met) or antagonize glucocorticoid receptors by injecting RU486 and test fear. (**M, N**) Conditioned and generalized fear. n=4-8 mice/group. (**O**) Neuron numbers in mice sacrificed immediately after behavior tests. n=3-4 mice/group. (**P**) Experimental design to interfere with glucocorticoid receptor (GR) expression using shRNA (shGR) and test fear. (**Q, R**) Conditioned and generalized fear. n=4-6 mice/group. (**S**) Neuron numbers in mice sacrificed immediately after behavior tests. n=3-4 mice/group.

We first gave strong footshock to activate CRH neurons in the paraventricular nucleus (PVN; CRH^PVN^), to trigger the release of CORT, and determined how long neuronal activity and CORT levels remained elevated. Co-staining of the PVN with CRH and c-fos, a marker for neuronal activity, revealed higher c-fos expression in CRH^PVN^ neurons 0.5 hour after shock. The increase in c-fos expression was sustained over the course of one day and decreased with time (**Fig. 4b,c**). Similarly, the level of plasma CORT was increased 0.5 hour after shock. At one day after shock, its level had decreased but remained higher in mice given shock than in no-shock mice (**Fig. 4d**).

We then investigated whether CORT, as the effector hormone of the HPA axis, is sufficient to induce the VGLUT3-to-GAD67 switch and produce generalized fear. We intraperitoneally injected a bolus of CORT (20 mg/kg) into mice and found that it was not sufficient to produce generalized fear (**Fig. 4e,f**). However, injection of CORT immediately before administering weak shock (0.5 mA) triggered both generalized fear and a VGLUT3-to-GAD67 switch in the 5-HT^lwDR^ neurons (**Fig. 4f,g**). Thus, a single injection of CORT combined with a weak shock was sufficient to establish generalized fear, although neither alone was sufficient (**Fig. 4g**).

To test whether activation of the HPA axis is required for generation of the neurotransmitter switch, we first suppressed the activity of its central regulator, the CRH^PVN^ neurons. We then examined the effect of suppressing their activity on the VGLUT3-to-GAD67 switch and on generalized fear. We injected CRH-Cre mice in the paraventricular nucleus with AAV-DIO-Kir2.1-zsGreen that expresses a Cre-dependent inwardly rectifying potassium channel (Kir2.1). After four weeks to allow full expression of Kir2.1, mice were given strong shock (**Fig. 4h**). Because Kir 2.1 was expressed before the footshock was delivered, it was expected to permanently suppress the activity of CRH^PVN^ neurons during and after the footshock. AAV-DIO-EGFP served as control. For another group of mice, we injected AAV-DIO-hM4D(Gi)-mCherry to express inhibitory chemogenetic receptors in the CRH^PVN^ neurons. Four weeks after the AAV injection when hM4D(Gi) was expressed in CRH^PVN^ neurons, we injected clozapine N-oxide (CNO), or saline as control, intraperitoneally into the mice expressing hM4D(Gi) and gave them strong shock 0.5 hours later (**Fig. 4h**). CNO was expected to block the activity of CRH^PVN^ neurons during the shock and for several hours afterward (*27*).

Generalized fear was reduced with expression of either Kir2.1 or activation of hM4D(Gi) by CNO (**Fig. 4i,j**). Both expression of Kir2.1 and activation of hM4D(Gi) by CNO reduced the number of 5-HT^lwDR^ neurons co-expressing GAD67 and increased the number of these neurons co-expressing VGLUT3 (**Fig. 4k**). Thus, both methods suppressed the VGLUT3-to-GAD67 switch (**Fig. 4h,k**). The expression of Kir2.1 produced greater suppression of fear and stronger suppression of the switch than activation of hM4D(Gi), likely because expression of Kir2.1 produced a stronger and more sustained suppression of activity than chemogenetic activation of the hM4D(Gi) receptor (**Supplementary Fig. 8**). Significantly, Kir2.1 reduced the number of neurons expressing GAD67 to control levels (cf **Fig. 4k** to **Fig. 1j and 1o** and **Fig. 5f**), consistent with selective effect on the neurons that would otherwise gain GAD67 in response to the strong shock. These results demonstrate that activation of the HPA axis is necessary to produce the transmitter switch.

**Fig. 5.**
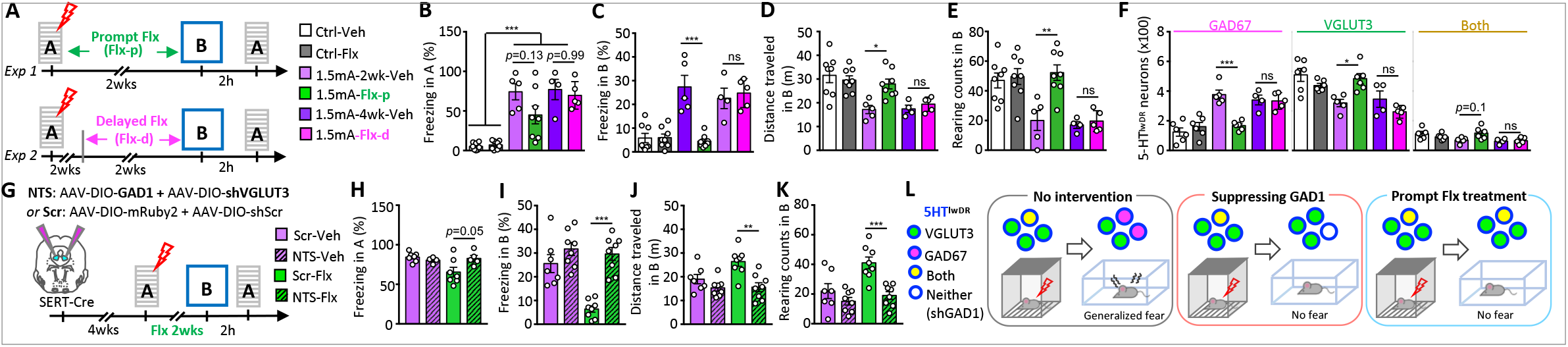
Prompt treatment with fluoxetine blocks generalized fear by interference with the co-transmitter switch in serotonergic neurons. (**A**) Experimental design for tests of fluoxetine treatment (Flx, 10 mg/kg/day). Veh, vehicle (drinking water). (**B, C**) Freezing in mice treated with Flx or Veh. (**D**, **E**) Exploration of mice in context B. (**B-F**) n=4-8 mice/group. (**F**) The number of 5-HT^lwDR^ neurons co-expressing GAD67, VGLUT3, or both. n=4-6 mice/group. (**G**) Experimental design to drive a VGLUT3-to GAD67 switch by co-injecting two AAVs and testing the effect of fluoxetine. (**H, I**) Freezing in mice expressing the AAV-DIO-shVGLUT3 plus AAV-DIO-GAD1 or expressing the two control vectors. Mice were given 5 1.5 mA shocks followed immediately with fluoxetine. (**J**, **K**) Exploration. (**H-K**) n=4-8 mice/group. (**L**) Left: Shock induces neurotransmitter switching from VGLUT3 to GAD67 and generalized fear. Middle: Suppressing GAD1 suppresses generalized fear. Right: Prompt treatment with fluoxetine blocks the VGLUT3-to-GAD67 switch and suppresses generalized fear.

CRH neurons in the CeA also regulate fear and project to the dorsal raphe (*6, 28, 29*). To test whether the activity of these CRH^CeA^ neurons contributes to the generation of the VGLUT3-to-GAD67 switch and sustained fear responses, we expressed AAV-DIO-Kir2.1-zsGreen in the CeA of CRH-Cre mice. Mice expressing Kir2.1 and mice expressing EGFP in the control virus both exhibited the transmitter switch and fear behavior (**Supplementary Fig. 9**), suggesting that CRH neurons in the CeA do not contribute to the VGLUT3-to-GAD67 switch nor, in agreement with an earlier report (*30*), to generalized fear.

We then tested whether blocking the synthesis of CORT or blocking glucocorticoid receptors (GR) suppressed the VGLUT3-to-GAD67 switch and blocked generalized fear. Prior to strong footshock, we intraperitoneally injected metyrapone (50 mg/kg) to block CORT synthesis or RU486 (40 mg/kg) to block GR (**Fig. 4l**). Metyrapone but not RU486 reduced conditioned fear in context A (**Fig. 4m**). Metyrapone blocked, and RU486 reduced, both generalized fear in context B and the transmitter switch (**Fig. 4n,o**). Additionally, we injected SERT-Cre mice with AAV-shGR (shRNA for GR) or AAV-shScr in the lwdr and compared freezing and expression of GAD67 and VGLUT3 (**Fig. 4p; Supplementary Fig. 6d**). We found that interfering with the GR in the 5-HT^lwDR^ blocked the footshock-induced switch, reduced conditioned fear in context A, and blocked generalized fear in context B (**Fig. 4q-s**). Taken together, these data demonstrate that activation of the HPA axis mediates the footshock-triggered VGLUT3-to-GAD67 switch and implicates the 5-HT^lwDR^ as a major target of CORT’s action.

### Immediate fluoxetine treatment blocks the glutamate-to-GABA switch and generalized fear

Delivery of fluoxetine has been found to reduce stress-induced conditioned fear (*31, 32*). To test whether fluoxetine similarly affects generalized fear, we gave mice fluoxetine in drinking water for two weeks either immediately after fear conditioning or beginning after a delay of two weeks (**Fig. 5a**). Fluoxetine consumption was similar in the two time periods (**Supplementary Fig. 10a**). At the end of two weeks of fluoxetine treatment, mice were tested in context B, one day later in context A, and then sacrificed (**Fig. 5a**). We found that prompt treatment, but not delayed treatment, partially reduced freezing in context A, fully blocked freezing in context B (**Fig. 5b,c**), and fully restored exploration behavior in context B (**Fig. 5d,e**) (see also *33*). The behavioral changes observed following immediate treatment with fluoxetine resembled the behavioral changes observed in shocked mice that expressed shGAD1 in the 5-HT^lwDR^ (**Fig. 2b,c**).

We then investigated how fluoxetine treatment affects neurotransmitter identity in the 5-HT^lwDR^. Prompt treatment with fluoxetine after strong shock blocked the gain of GAD67 and loss of VGLUT3 in 5-HT^lwDR^ neurons (**Fig. 5f**). Delayed treatment did not prevent the VGLUT3-to-GAD67 switch. These results demonstrate a time-sensitive period in which fluoxetine can block the VGLUT3-to-GAD67 switch in 5-HT^lwDR^ neurons and block acquisition of both conditioned and generalized fear.

Ordinarily, the number of 5-HT^lwDR^ neurons co-expressing GAD67 is inversely correlated with the number of 5-HT^lwDR^ neurons co-expressing VGLUT3 and positively correlated with the extent of generalized fear. To determine whether the VGLUT3-to-GAD67 switch is necessary for fluoxetine to block generalized fear, we drove an exogenous VGLUT3-to-GAD67 switch in 5-HT^lwDR^ neurons by co-injecting AAV-DIO-shVGLUT3-EGFP and AAV-DIO-GAD1-P2A-mRuby2 into the 5-HT^lwDR^ of SERT-Cre mice (**Fig. 5g**). Co-expression of the two AAVs in 5-HT^lwDR^ neurons was confirmed by the co-localization of EGFP, mRuby2 and 5-HT in 5-HT^lwDR^ (**Supplementary Fig. 6a,e,f**). Mice injected with AAV-DIO-shScr-EGFP and AAV-DIO-mRuby2 served as controls. Comparing the level of VGLUT3 in shVGLUT3-expressing 5-HT^lwDR^ neurons with the level in neurons expressing shScr indicated that the efficiency of knocking down VGLUT3 by AAV-DIO-shVGLUT3-EGFP was 79% (**Supplementary Fig. 6e**). Knockdown of VGLUT3, together with exogenous expression of GAD67, fully prevented the block of generalized fear produced by fluoxetine (**Fig. 5i-l**). The consumption of fluoxetine was similar for mice expressing shVGLUT3 and GAD1 and mice expressing shScr and mRuby2 (**Supplementary Fig. 10b**). These results suggest that fluoxetine suppresses generalized fear by interfering with the VGLUT3-to-GAD67 switch (**Fig. 5l**).

Because a VGLUT3-to-GAD67 switch prevents fluoxetine from reducing generalized fear, we asked whether driving the switch with viral tools would be sufficient to produce generalized fear. Accordingly, we co-injected AAV-DIO-shVGLUT3-EGFP and AAV-DIO-GAD1-P2A-mRuby2 into the lwDR of SERT-Cre mice (**Supplementary Fig. 11a**). Mice injected with AAV-DIO-shScr-EGFP and AAV-DIO-mRuby2 served as controls. When the transmitter switch was driven by viral tools, freezing was greater in mice expressing shVGLUT3 and GAD1 than mice given no shock (9% vs 3%) (**Supplementary Fig. 11b**), but still less than mice given strong shock (29%; **Fig. 1c**). Rearing was decreased but distance traveled and number of fecal boli were not changed by the expression of shVGLUT3 and GAD1 (**Supplementary Fig. 11c-e**). These data demonstrate that driving a VGLUT3-to-GAD67 switch is sufficient to produce a low level of generalized fear. The difference between the effect of driving the switch with viral tools and with strong footshock acknowledges the additional contributions to the generation of strong fear behavior after shock, such as elevation of CORT.

## Discussion

Our results show that acute stress triggers a sustained co-transmitter switch from glutamate/VGLUT3 to GABA/GAD67 in a population of 5-HT^lwDR^ neurons innervating the CeA and LHA. Overriding the gain of GABA by suppressing the expression of GAD1 blocks generalized fear. Initiation of the transmitter switch is achieved by activation of the CRH-CORT-GR pathway. Blocking this pathway or immediate treatment with fluoxetine prevents the VGLUT3-to-GAD67 switch and blocks generalized fear. It will be interesting to learn whether the VGLUT3-to-GAD67 switch in the 5-HT^lwDR^ occurs in response to other forms of physical pain or psychological stress.

The relationship between neurotransmitter co-expression and switching has been of substantial interest (*34, 35*). Neurons that co-express two transmitters can lose one transmitter while retaining the other (*36–38*). We find that one co-transmitter marker (VGLUT3) can be replaced by another (GAD67), while a third transmitter (5-HT) does not change, demonstrating the co-occurrence of transmitter co-expression and transmitter switching. Transmitter changes in neurons co-expressing multiple transmitters have been linked to changes in motor function and therapies for Parkinson’s disease (*34, 39–41*).

We demonstrate that generalized and conditioned fear differ in induction threshold to shock (*42*), in time course (*12*), and in dependence on the VGLUT3-to-GAD67 switch in the 5-HT^lwDR^. A single shock of 0.5 mA was sufficient to produce conditioned fear but produced neither the transmitter switch nor generalized fear. Conditioned fear was observed 1 day after the shock, but the transmitter switch and generalized fear were absent at 1 day and present at 3 days after the shock. Generalized fear showed a close correlation with and a tight dependence on the switch. When the switch was blocked, generalized fear disappeared. The switch does not affect early expression of conditioned fear since conditioned fear is present before the transmitter switch has occurred. However, blocking the switch reduces conditioned fear at a later time point - 14 days after footshock - suggesting that the switch contributes to the later phase of conditioned fear.

Sustained conditioned fear and generalized fear are both controlled by neurons in the medial prefrontal cortex (mPFC) (*43, 44*) and amygdala (*45, 46*), consistent with convergence of the pathways regulating the two forms of fear. We find that sustained generalized fear is regulated by the CeA and LHA, which are downstream targets for switching neurons in the 5-HT^lwDR^. Infusion of agonists for GABA receptors in a subregion of the CeA induces unconditioned freezing behavior (*47*), suggesting that the 5-HT^lwDR^ neurons that have switched to co-expressing GAD67/GABA may release GABA into the CeA and trigger generalized fear. The neurons in the CeA and LHA that are innervated by switching neurons remain unclear, but may include CeA neurons expressing protein kinase C δ (*48*) or somatostatin (*49*), and glutamatergic neurons in the LHA (*50*).

We show that corticosterone and its receptors are required for the switch from VGLUT3 to GAD67/GABA in 5-HT^lwDR^ neurons. These results are consistent with observations that injection of corticosterone increases GABA release and reduces glutamate release in the adult rodent amygdala (*51*) and upregulates GAD67 mRNA in the hippocampus (*52*). Chronic treatment with antidepressants inhibits glucocorticoid receptor-mediated gene transcription (*53, 54*) and reverses corticosterone-induced anxiety/depression-like behavior (*55*). Thus, prompt fluoxetine may block the transmitter switch in the 5-HT^lwDR^ by blocking glucocorticoid receptor-mediated gene transcription in 5-HT^lwDR^ neurons. We found that treatment with fluoxetine two weeks after footshock did not block transmitter switching and acquisition of generalized fear. The short time window for effective therapy may explain why PTSD patients with long histories of this disorder are less responsive to SSRIs (*56*).

## Materials and Methods

### Mice

All animal procedures were carried out in accordance with NIH guidelines and approved by the University of California, San Diego Institutional Animal Care and Use Committee. C57BL/6J (JAX#000664), VGLUT3-Cre (JAX#018147), Rosa26-CAG-LSL-H2B-mCherry (JAX#023139), SERT-IRES-Cre (JAX#014554), and CRH-IRES-Cre (#012704) mice were obtained from Jackson Laboratories. Animals were maintained on a 12 h:12 h light:dark cycle (light on: 8:00 am–8:00 pm) with food and water ad libitum and single housed one week before undergoing experiments. Vivarium temperature was maintained between 18 and 23 °C and humidity between 40% and 60%. VGLUT3-Cre male mice and Rosa26-CAG-LSL-H2B-mCherry homozygous female mice were used to generate VGLUT3-Cre::mCherry-H2B double transgenic mice. SERT-IRES-Cre and CRH-IRES-Cre colonies were maintained by breeding homozygous transgenic males with female wild-type C57BL/6J mice. Heterozygous VGLUT3-Cre::mCherry-H2B, SERT-IRES-Cre, CRH-IRES-Cre offspring were used in the study. Experiments were done on males unless otherwise specified. All experiments were performed on 2-to 4-month-old male and female mice.

### Fear conditioning

We used contextual fear as the CS associated with the US; no other CS, such as light or tone, was introduced. Before fear conditioning, we habituated mice to the room for 1 h and transferred them into the chamber (CS, SuperFlex Shuttle Box, Omnitech Electronics Inc.) one at a time for fear conditioning. Each mouse was habituated to the chamber for 2 min and footshocks (US) were delivered through a metal grid on the floor. For weak shock experiments, each mouse received one shock at an intensity of 0.5 mA and a duration of 1 s. For intense shock experiments, each mouse received five shocks, at an intensity of 1.5 mA and a duration of 2 s. The intervals between shocks varied randomly from 40 to 60 s. For both paradigms, mice were kept in the chamber for one minute after the last shock to reinforce fear memory associated with the chamber. The chamber was cleaned with 70% ethanol before and after each mouse.

### Tests of conditioned and generalized fear

After fear conditioning, re-exposure of an animal to the CS (CS+) without the US enables measurements of conditioned fear (*57*). Exposure to different environments (CS-) allows evaluation of generalized fear (*13–15*). Two weeks after footshock, we measured how mice react to both the conditioned context (CS+; context A) and to two different contexts (CS-; context B or C). The three contexts had different shapes, sizes, textures and colors. Context A was a 20cmx20cmx30cm (LxWxH) chamber with one black plastic, one metal and two transparent walls, a stainless-steel grid floor and a metal lid; it was maintained under normal room light (100 lux), and cleaned with 70% ethanol before and after testing each mouse. Context B was a 50cmx50cmx30cm (LxWxH) chamber with four white plastic walls and floor, no lid, maintained under bright room light (250 lux), and cleaned with lemon scent disinfecting wipes and dry paper towels before and after testing each mouse. Context C was a black plus-maze made of two open (50cmx5cmx1cm, LxWxH) and two closed arms (50cmx5cmx20cm, LxWxH), with no lid, maintained under dim light (20 lux), and cleaned with 10% bleach and dry paper towels before and after testing each mouse.

### Acoustic startle response and prepulse inhibition

SR-LAB startle system (San Diego Instruments) was used to measure acoustic startle response (ASR) and prepulse inhibition (PPI). Before each experiment, the system was calibrated with a digital sound meter. Mice were acclimated to the testing room for 1 h before the test and then gently transferred into an animal enclosure setup inside the cabinet of the startle system. The animal enclosure setup was coupled to an accelerometer sensor that digitized the pressure generated by startled mice. Data were collected using SR-LAB software (San Diego Instruments) and ASR amplitude was analyzed using SR-Analysis software (San Diego Instruments). Each session began with 5 min of habituation, followed by 25 trials of ASR and then 25 trials of PPI. The duration of the entire session was 20±3 min. During the habituation phase, the chamber was buffered with 40 dB background noise. Each ASR trial consisted of a white noise stimulus with an intensity of 110 dB and duration of 30 ms. Each PPI trial consisted of a prepulse white noise stimulus with an intensity of 70 dB and duration of 30 ms, followed by an interval of 90 ms, and then a white noise stimulus with an intensity of 110 dB and duration of 30 ms. Trials were separated by random intervals (ITIs) ranging from 10-25 s. The resulting startle reflexes of mice in the startle chamber were recorded during the first 500 ms starting from the onset of each acoustic stimulus. ASR was defined as the amplitude (voltage) of the largest response that occurred within 200 ms after the onset of the startle stimulus and was quantified as the mean value of 25 startle trials. The PPI was calculated as

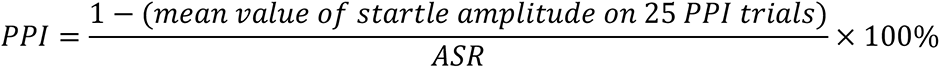

### Corticosterone measurement

Whole submandibular (chin) blood of non-anesthetized mice was collected using lancets and Eppendorf tubes. Blood samples were collected between 9:00 am and 12:00 pm. Plasma was separated by centrifugation at 10,000 g at 4°C for five minutes and stored at −80 °C until assayed. Corticosterone levels were detected using the Corticosterone EIA Kit (K014-H5; Arbor Assays) according to the manufacturer’s protocol.

### Drug administration

Mice were intraperitoneally injected with 50 mg/kg metyrapone (3292; R&D) to block CORT synthesis. To block glucocorticoid receptors, mice were intraperitoneally injected with 40 mg/kg RU-486 (also referred as mifepristone; M8046; Sigma-Aldrich). Metyrapone and RU-486 were freshly dissolved in DMSO and then diluted 1:100 in pure corn oil. Injection with 1% DMSO/corn oil served as a vehicle control. Mice were returned to their home cages after i.p. injection and given footshock 40 min later.

For acute CORT exposure, mice were intraperitoneally injected with 10 mg/kg corticosterone 2-hydroxypropyl-β-cyclodextrin complex (CORT-HBC; C174; Sigma-Aldrich) and returned to their home cage. 2-hydroxypropyl-β-cyclodextrin complex (HBC; H107; Sigma-Aldrich) was used as vehicle control. Mice were given footshock immediately after i.p. injection of CORT-HBC or HBC and then returned to their home cages.

For DREADD-based chemogenetic inactivation, mice expressing hM4D(Gi) were intraperitoneally injected with 2 mg/kg CNO (89160; VWR International, LLC) and returned to their home cages. Saline was used as control. Mice were given footshock thirty minutes after the i.p. injection.

For fluoxetine delivery, mice were provided with fluoxetine in drinking water. Fluoxetine solution was made fresh each day at a concentration of 1 mg/mL to reach a daily intake of approximately 10 mg/kg per day (*31*). Fluoxetine solution was provided to mice in polypropylene light-protected tubes that were weighed daily to calculate the mount of drug intake. For control mice, the same type of tube provided drinking water and water was changed daily to introduce similar disturbances to control and to fluoxetine-treated mice.

### Histology and immunocytochemistry of mouse brains

Mice were perfused transcardially with phosphate-buffered saline (PBS) followed by 4% paraformaldehyde (PFA) in PBS. Brains were dissected and post-fixed in 4% PFA for 16 to 24 h at 4 °C, washed in PBS for 1 min and transferred to 30% sucrose in PBS for 2 days at 4 °C. Forty µm coronal sections were cut on a microtome (Leica SM2010R) and stored in cryoprotectant at -20°C. Before immunostaining, sections were washed in PBS at 4 °C overnight and then permeabilized and blocked in 24-well culture plates for 2 h in a blocking solution (5% normal horse serum, 0.3% Triton X-100 in PBS) at 22–24 °C. Primary and secondary antibodies were diluted in the blocking solution. Incubation with primary antibodies was performed for 48 h on a rotator at 4 °C. After washing in PBS (3 times, 15 min each), secondary antibodies were added for 2 h at 22–24 °C. For immunofluorescence, sections were mounted with Fluoromount-G (Southern Biotech) or ProLong Gold Antifade Mountant (Life Technologies) containing DRAQ-5 (Thermo Fisher, 62251, 1:1000 dilution, when nuclear staining was needed) after washes in PBS (three times, 15 min each).

Primary antibodies used in this study were goat anti-serotonin (ImmunoStar, 20079, 1:1000), rabbit-anti-VGLUT3 (Synaptic Systems, 135203, 1:500), guinea pig-anti-VGLUT3 (Synaptic Systems, 135204, 1:500), mouse-anti-GAD67 (Millipore, MAB5406, 1:500), mouse-anti-cFos (Abcam, ab208942, 1:500), guinea pig-anti-VGAT (Synaptic Systems, 131004, 1:1000), rabbit-anti-GABA (Sigma-Aldrich, A2052, 1:1000) rabbit-anti-GFP (Thermo Fisher, A11122, 1:1000), chicken anti-GFP (Abcam, ab13970, 1:1000), rabbit-anti-zsGreen (Takara, 632474, 1:500), chicken-anti-mCherry (Abcam, ab125096, 1:500), goat-anti-doublecortin (Santa Cruz, sc-8066, 1:300) and rabbit-anti-Ki67 (Cell Signaling, 9129, 1:300). Secondary antibodies for immunofluorescence were from Jackson ImmunoResearch Labs and used at a concentration of 1:600: Alexa Fluor-488 donkey-anti-rabbit (705-545-003), Alexa Fluor-488 donkey-anti-guinea pig (706-545-148), Alexa Fluor-488 donkey-anti-mouse (715-545-150), Alexa Fluor-488 donkey-anti-goat (705-545-147), Alexa Fluor-594 donkey-anti-goat (705-585-147), Alexa Fluor-594 donkey-anti-mouse (715-585-150), Alexa Fluor-594 donkey-anti-chicken (703-585-155), Alexa Fluor-647 donkey-anti-goat (705-605-147), Alexa Fluor-647 donkey-anti-rabbit (711-605-152), Alexa Fluor-647 donkey-anti-guinea pig (706-545-148) and Alexa Fluor-680 donkey-anti-goat (705-625-147).

### Histology and immunocytochemistry of human brains

Postmortem midbrain tissues were provided by NIH NeuroBioBank and shipped in cryoprotectant. Upon arrival, tissues were kept in cryoprotectant at -20 °C, cut into 2 cm x 2 cm x 1 cm (LxWxH) tissue blocks using lancets and sectioned into 80 µm coronal sections on a microtome (Leica SM2010R). Brain sections were stored in cryoprotectant at -20 °C and washed in PBS at 4 °C overnight. For immunostaining, brain sections were first mounted on coded slides and let air-dry. Sections were then incubated in 1x Universal HIER antigen retrieval solution (Abcam, ab208572) at 95 °C, followed by cooling down to the room temperature and washing in diH_2_0 (twice, 1 min each). Sections were permeabilized and blocked for 3 h in a blocking solution (5% normal horse serum, 0.3% Triton X-100 in PBS) at 22–24 °C. Primary and secondary antibodies were prepared in the blocking solution. Incubation with primary antibodies was performed for 4 d on a rotator at 4 °C to allow full penetration and incubation. After washing in PBS (3 times, 15 min each), secondary antibodies were added for 3 h at 22–24 °C. After washes in PBS (three times, 15 min each), sections were incubated in TrueBlack® Lipofuscin autofluorescence quencher (Biotium, 23007) for 2 min, rinsed in PBS for 1 min, and mounted with ProLong Gold Antifade Mountant (Life Technologies). Primary antibodies used in this study were goat anti-TPH2 (Abcam, ab121013, 1:500), mouse-anti-GAD67 (Abcam, ab26116, 1:200), and rabbit-anti-VGLUT3 (Millipore, AB5421-II: 1:200). Secondary antibodies for immunofluorescence were from Jackson ImmunoResearch Labs and used at a concentration of 1:600: Alexa Fluor-488 donkey-anti-rabbit (705-545-003, Alexa Fluor-594 donkey-anti-mouse (715-585-150) and Alexa Fluor-647 donkey-anti-goat (705-605-147).

### TUNEL assay

To detect in situ apoptosis, 40 µm cryosections were re-fixed with 1% PFA for 20 min at 22–24 °C and rinsed with PBS (three times, 5 min each). Sections were then permeabilized in 0.1% sodium citrate and 1% Triton X-100 for 1 h at 22–24 °C. After rinsing in PBS (three times, 5 min each), sections were incubated with In Situ Cell Death Detection (TUNEL) Kit with TMR Red (Roche, 12156792910) according to the vendor’s instructions: incubated in a mixture of 25 μL of terminal-deoxynucleotidyl transferase solution and 225 μL of label solution. Incubation was performed in a humidified chamber for 3 h at 37 °C in the dark. Sections were rinsed and mounted with Fluoromount containing DRAQ-5 (1:1000). For a positive control, sections were treated with DNase I (10 U/mL, New England Biolabs, M0303S) for 1 h at 37 °C and rinsed in PBS (three times, 5 min each), followed by incubation with the TUNEL mixture.

### Imaging and data analysis

Fluorescent images were acquired with Leica Stellaris 5, Leica SP8 or Leica SP5 confocal microscopes. 20x/0.75 dry and 25x/0.95 water-immersion objectives were used to image cell bodies. A 63x/0.90 water-immersion objective at 2x zoom magnification was used for synaptophysin-EGFP signals in nerve terminals. All images were acquired at a z resolution of 1 μm. Colocalization was quantified using Image J with the JACoP plugin. To delimit the lwDR in sections stained for Ki67/DCX (apoptosis and neurogenesis, Supplementary Fig.3), an adjacent section was stained for 5-HT to define the boundaries of the lwDR. Leica Application Suite X software was used for fluorescent cell counting and all sections within confocal stacks were examined without maximal projection. Images were analyzed exhaustively or by sampling. Exhaustive analysis entailed scoring all neurons in every sixth section through the entire brain region of interest and reporting the number of cells for the number of mice examined. Sampling involved scoring and reporting a specific number of cells from a specified number of sections from a particular number of mice. In these cases, exact cell and/or section numbers are shown in the figure legends.

### Viral constructs and injection

Eight-week-old male mice were anesthetized with a mixture of 120 mg/kg ketamine and 16 mg/kg xylazine and head-fixed on a stereotaxic apparatus (David Kopf Instruments Model 1900) for all stereotaxic surgeries. The lwDR, PVN, LHA and CeA were targeted bilaterally. Stereotaxic coronates were: lwDR (AP, −4.60 mm from bregma; ML, ±0.7 mm; DV, −2.6 mm from the dura), PVN (AP, -0.80 mm; ML, ± 0.30 mm; DV, -4.70 mm), LHA (AP −1.2 mm; ML ±1 mm; and DV from the dura, −5.1 mm), and CeA (AP, −1.2 mm; ML, ±2.25 mm; and DV, −4.65 mm). A total of 300 nL of the Cre-on viral vectors were injected into each lwDR of SERT-Cre mice at a rate of 100 nL/min using a syringe pump (PHD Ultra™, Harvard apparatus, no. 70-3007) installed with a microliter syringe (Hampton, no.1482452A) and capillary glass pipettes with filament (Warner Instruments, no. G150TF-4). Pipettes were left in place for 5 min after injection. Titers of recombinant AAV vectors ranged from 7.1 × 10^12^ to 2.2 × 10^13^ viral particles/ml, based on quantitative PCR analysis. AAV8-phSyn1(S)-FLEX-tdTomato-T2A-SypEGFP-WPRE was from the Salk Viral Vector Core (*17, 22*). AAV-DJ-CMV-DIO-Kir2.1-zsGreen, AAV-DJ-CMV-DIO-EGFP, and AAV-DJ-CMV-DIO-hM4Di-mCherry were from the Stanford Gene Viral Vector Core (*17, 38, 58, 59*). AAV8-hSyn-DIO-GAD1-P2A-mRuby2, AAV-hSyn-DIO-VGLUT3-P2A-mRuby2 and AAV8-hSyn-DIO-mRuby2 were constructed in the Cleveland lab. The P2A self-cleavage sequence located between GAD1 or VGLUT3 and mRuby2 separates the two proteins and prevents potential dysfunction of GAD1 or VGLUT3 by fusion with mRuby2. Vector Biolabs produced AAV8-CAG-DIO-shRNAmir-mGAD1-EGFP (*17, 60*), AAV8-CAG-DIO-shRNAmir-mVGLUT3-EGFP, AAV8-CAG-DIO-shRNAmir-Scramble-EGFP, AAV8-CAG-shRNAmir-GR-EGFP, and AAV8-CAG-shRNAmir-Scramble-EGFP.

The shRNA sequence for mouse GAD1 is 5′-GTCTACAGTCAACCAGGATCTGGTTTTGGCCACTGACTGACCAGATCCTTTGACTGTAGA-3′.

The shRNA sequence for mouse VGLUT3 is 5’-GCTGAAAGTAAGCTGGCTGACTTATGTTTTGGCCACTGACTGACATAAGTCACAGCTTACTTT CAG-3’.

The shRNA sequence for GR is 5’-CCGGTGAGATTCGAATGACTTATATCTCGAGATATAAGTCATTCGAATCTCATTTTT-3’.

### Tract tracing

For anterograde tracing, 300 nL of AAV8-phSyn1(S)-FLEX-tdTomato-T2A-SypEGFP-WPRE were bilaterally injected into the lwDR of SERT-Cre mice and EGFP+ synaptic terminals were imaged by Leica confocal microscope. For retrograde tracing, a volume of 100 nL of FluoroGold (Biotium, 80023) was stereotaxically injected at 100 nL/min into the LHA or CeA. Pipettes were left in place for 10 min after injection. Two weeks after injection, mice were sacrificed for brain examination or foot-shocked to examine stress-induced loss of VGLUT3 and gain of GAD67 in 5-HT^lwDR^ neurons that contained FluoroGold.

### Statistics and reproducibility

For comparisons between two groups, data were analyzed by Welch’s t-test for normally distributed data and the Mann–Whitney U test for data not normally distributed, using Graph Pad Prism 7. For repeated samples, data were analyzed by paired t-test. Correlation was analyzed by the Pearson correlation using Graph Pad Prism 7. All tests were two-tailed. Statistical analyzes of the data were performed using Prism 7 software for the number of animals for each experiment indicated in the figure legends. Means and SEMs are reported for all experiments. For comparisons between multiple groups, ANOVA followed by Tukey’s test was used for normally distributed data. The nonparametric Kruskal–Wallis test followed by Dunn’s correction was used for data that were not normally distributed.

## Acknowledgments

We thank all members of the Spitzer lab for discussions and useful critiques, Alex Glavis-Bloom for technical support, and Romona Dong and Rahul Sharmarayar for assistance with experimental procedures. We thank D.K. Berg and L.R. Squire for comments on the manuscript.

## Author contributions

H.-q.L. and N.C.S. conceived the study, designed the experiments, interpreted the data and wrote the manuscript. H.-q.L. performed most of the experiments and data analysis. W.J. performed cell birth/death experiments, assisted with studies of fluoxetine, and contributed to data analysis. L.L performed CORT detection experiments and assisted with studies involving the HPA axis. V.G. assisted with studies of fluoxetine and AAV injection. C.C. designed, constructed and validated AAV tools. M.P. and S.G. contributed to experimental design and data interpretation.

## Data and materials availability

All data are available in the manuscript or the supplementary materials.

